# Targeting intracellular of populations *Pseudomonas aeruginosa* with peptide-mimetic therapies: individual efficacy and synergistic rescue of obsolete antibiotics

**DOI:** 10.1101/2025.08.26.672426

**Authors:** Melanie L. Burger, Rose Doyle, Jennifer S. Lin, Annelise E. Barron, Madeleine G. Moule

**Affiliations:** Institute of Immunology and Infection Research, School of Biological Sciences, University of Edinburgh, Edinburgh, UK; Department of Bioengineering, School of Medicine & School of Engineering, Stanford University, Stanford, California 94305, United States

## Abstract

*Pseudomonas aeruginosa* is a leading cause of human infections, with current treatment options severely limited by high levels of antimicrobial resistance. Historically considered to be an extracellular pathogen, recent evidence has emerged that *P. aeruginosa* is able to survive and replicate within human cells. These intracellular niches present an additional clinical challenge and may serve as bacterial reservoirs associated with chronic infections that are particularly difficult to eradicate. Here we describe the application of a novel peptide-based therapeutic against recalcitrant populations of bacteria residing within lung epithelial cells. This antimicrobial “peptoid” is able to target intracellular bacteria without harming host cells. In addition, we have shown that peptoid TM5 exhibits synergy with three antibiotics that otherwise have low efficacy against *P. aeruginosa*, effectively rescuing drugs that have become clinically obsolete. These synergistic combination therapies are also capable of reducing intracellular bacterial reservoirs, opening the door for potential new strategies against chronic *P. aeruginosa* infections.

## Introduction

The overuse of antibiotics has led to multidrug resistant (MDR) bacteria becoming one of the biggest threats to global health. In 2024 the WHO published its updated Bacterial Priority Pathogens List (BPPL) which highlighted the global need for novel antibiotics against antibiotic-resistant pathogens, such as *Salmonella, Shigella*, *Neisseria gonorrhoeae*, *Pseudomonas aeruginosa*, and *Staphylococcus aureus*. These pathogens are responsible for the majority of MDR nosocomial infections and are able to evolve resistance mechanisms faster than new therapeutics can be produced^1, 2^.

The gram-negative, opportunistic pathogen *P. aeruginosa* possesses many intrinsic, adaptive and acquired antibiotic resistance mechanisms. Carbapenem resistance in particular is highly correlated with increased mortality and significantly more severe disease^3, 4^. A 2019 report estimates that globally *P. aeruginosa* causes > 500,00 deaths each year and that 300,000 of these deaths are MDR related. MDR *P. aeruginosa* causes 7% of all healthcare infections, as the pathogen can be found in contaminated sinks, toilets and medical equipment^5^. This exposes vulnerable patients, such as those who are elderly or immunocompromised, to contracting *P. aeruginosa* infections, which can lead to pneumonia, urinary tract infections and bacteraemia. *P. aeruginosa* poses a particularly serious threat for cystic fibrosis (CF) patients and is responsible for the majority of the mortality of these patients, mainly due to its ability to cause chronic infections^6^.

CF is a genetic disorder caused by multiple mutations in the gene encoding the cystic fibrosis transmembrane conductance regulator (CFTR) protein^7^. Dysfunctional CFTR leads to thickening of secretions and mucus build-up in the lungs and digestive tract. Most patients’ lungs will become colonized with bacteria at an early age and this mucus build-up provides an excellent environment for biofilms to form, which in turn facilitates the establishment of chronic infections and further emergence of antimicrobial resistance^8–10^ These factors, combined with the lack of development of novel therapeutics, make *P. aeruginosa* a serious health risk for CF patients^11^.

Historically, *P. aeruginosa* has been classed as an extracellular pathogen, although it was first observed to be internalized by lung epithelial cells in the 1990s^12^. Only recently has the pathogen’s intracellular life cycle stage gained more interest as more evidence has emerged that *P. aeruginosa* is able to survive and even replicate inside host cells, creating a bacterial reservoir that has been proposed to contribute to chronic infections such as those often found in CF patients^13–18^. These previously understudied intracellular populations of bacteria are likely be a significant barrier to the treatment of chronic infections as intracellular bacteria are likely to be protected from elimination by many types of antibiotics^19^. In addition, there is evidence that the CFTR plays a role in the internalization of *P. aeruginosa*, and that CFTR mutations may lead to increased intracellular bacterial growth, making the treatment of intracellular *P. aeruginosa* a particularly relevant issue for CF patients ^20, 21^.

Tobramycin, the gold standard antibiotic used to treat *P. aeruginosa* infections is an aminoglycoside and therefore not host cell permeable, making it unable to eliminate intracellular bacteria^22, 23^. Data has shown that another first-line antibiotic, ciprofloxacin, is able to treat intracellular *P. aeruginosa* but is not effective against all infections due to high levels of resistance and its association with potentially severe side effects^24, 25^. Due to these limitations, targeting not only extracellular, but also intracellular *P. aeruginosa* should be taken into consideration when choosing treatment regimens and developing new ones.

Antimicrobial peptides (AMPs) have long been seen as a promising alternative to conventional antibiotics. These small molecules naturally produced by the innate immune system of nearly all organisms display an extensive spectrum of antimicrobial activity^26^. AMPs are also associated with a decreased risk of resistance evolution as they can kill bacteria through different mechanisms and can stimulate the host immune system^25, 27^. Despite their advantages, the clinical use of AMPs has been limited, often due to their instability and potential cytotoxicity to host cells^28^.

We previously developed peptidomimetic derivates of the human AMP LL-37^29–31^. The bioavailability of LL-37 is problematically low due to sensitivity to degradation by proteases, and failure to efficiently clear bacterial infections has hindered its clinical development^32^. Our synthetic derivatives, known as peptoids or *N*-substituted glycine oligomer peptidomimetics, are resistant to proteolysis due to structural modifications. The active side chains of peptoids are linked to backbone amide nitrogen instead of α-carbon and can therefore no longer form hydrogen bonds within their main chain^29, 33, 34^. We recently showed that our antimicrobial peptoids are effective at clearing *P. aeruginosa* infections in a respiratory infection mouse model and are less cytotoxic in *in vivo* and *in vivo-*mimicking conditions. In particular, peptoid TM5 was shown to be well tolerated at high concentrations and efficient at clearing *P. aeruginosa* infections^31^. TM5 has also exhibited activity against *P. aeruginosa* biofilms and its persister cells^35^. In this study, we sought to further optimize TM5 as a potential therapeutic by exploring the potential of this peptide to eliminate intracellular populations of bacteria both alone and in combination with conventional antibiotics.

Many AMPs have been shown to work excellently in combination with other peptides, conventional antibiotics or various nanoparticles^36^. Simultaneous administration of some drug pairings can lead to a synergistic effect, in which the combination of drugs has a stronger effect than that of a single drug. Synergistic applications with antibiotics could potentially diminish some of the challenges facing clinical applications of AMPs. Minimal inhibitory concentrations and toxicity can be significantly reduced when AMPs are used in synergy with antibiotics because the concentrations of each therapeutic can be decreased^37, 38^. Antibiotics that are no longer in use can even be “recycled” when AMPs are applied at the same time, as some AMPs can disrupt bacterial membranes allowing increased penetration of the antibiotic ^39^. Here we provide evidence for potential synergistic applications of TM5 in combination therapy approaches against *P. aeruginosa.* Furthermore, we demonstrate the ability of TM5 alone and as part of synergistic combinations therapies to treat intracellular *P. aeruginosa* infections. Our results suggest that TM5 may have broader applications against chronic MDR *P. aeruginosa* infections and pave the way for the development of new clinical therapy regimens.

## Results

### Establishment of models of intracellular *P. aeruginosa* infection

We previously showed that peptoid TM5 is highly effective both *in vitro* and i*n vivo* against *P. aeruginosa*^31^. Using standard broth microdilution assays, we determined that TM5 has a minimum inhibitory concentration (MIC) of 8 μg/mL in both laboratory and clinical strains of *P.* aeruginosa, and a minimum bactericidal concentration of 32 μg/mL for all strains tested (Figure 1). To determine if TM5 has further applicability against chronic CF lung infections, we next sought to evaluate whether the peptoid is effective against intracellular populations of bacteria using the human lung epithelial cell line Calu-3.

**Figure 1.**
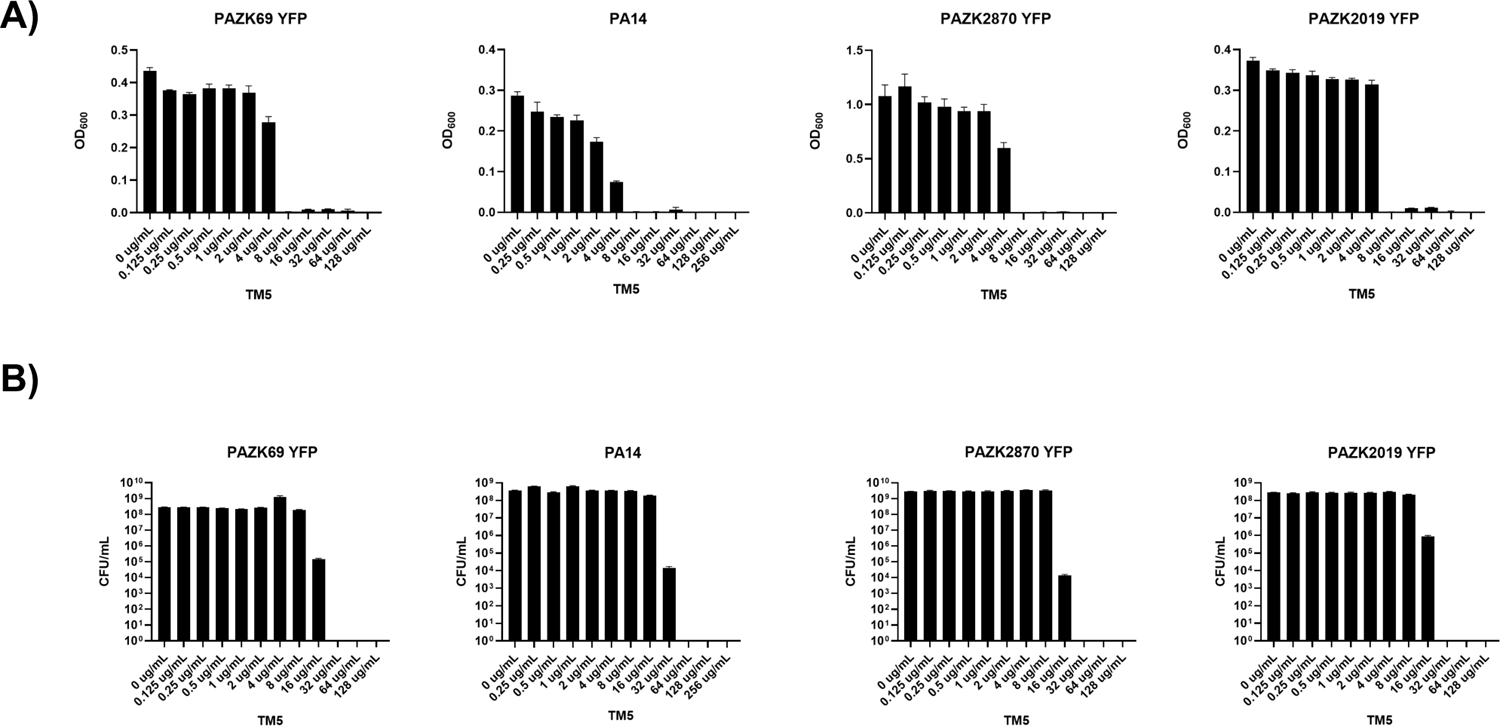
Growth inhibition (MIC) and bactericidal (MBC) concentrations of peptoid TM5 a) Minimum inhibitory concentrations (MIC) of peptoid TM5 against *P. aeruginosa* PAO1 as well as the clinical isolates *P. aeruginosa* PAZK2019, PAZK69, and PAZK2879. b) Minimum bactericidal concentrations (MBC) for each strain as measured by bacterial CFU assay

To confirm that we were in fact treating intracellular bacteria, we first validated our infection model to demonstrate the presence of intracellular *P. aeruginosa* within Calu-3 cells using a standard antibiotic protection assay to kill extracellular bacteria^40^. Lung epithelial cells were infected with *P. aeruginosa* PA01, and following a 1-hour incubation the cell-impermeable antibiotic amikacin was applied to eliminate all extracellular bacteria. The cells were then washed with PBS to remove any remaining antibiotic, and the supernatant, wash buffer and cellular lysates were plated onto LB agar plate. Following incubation, bacterial colony forming units (CFUs) were enumerated to ensure the presence of intracellular and eradication of extracellular bacteria (Figure 2a). Further validation was performed using confocal microscopy of infected Calu-3 cells and A549 lung epithelial cells to confirm the presence of intracellular *P. aeruginosa* (Fig 2b,c).

**Figure 2.**
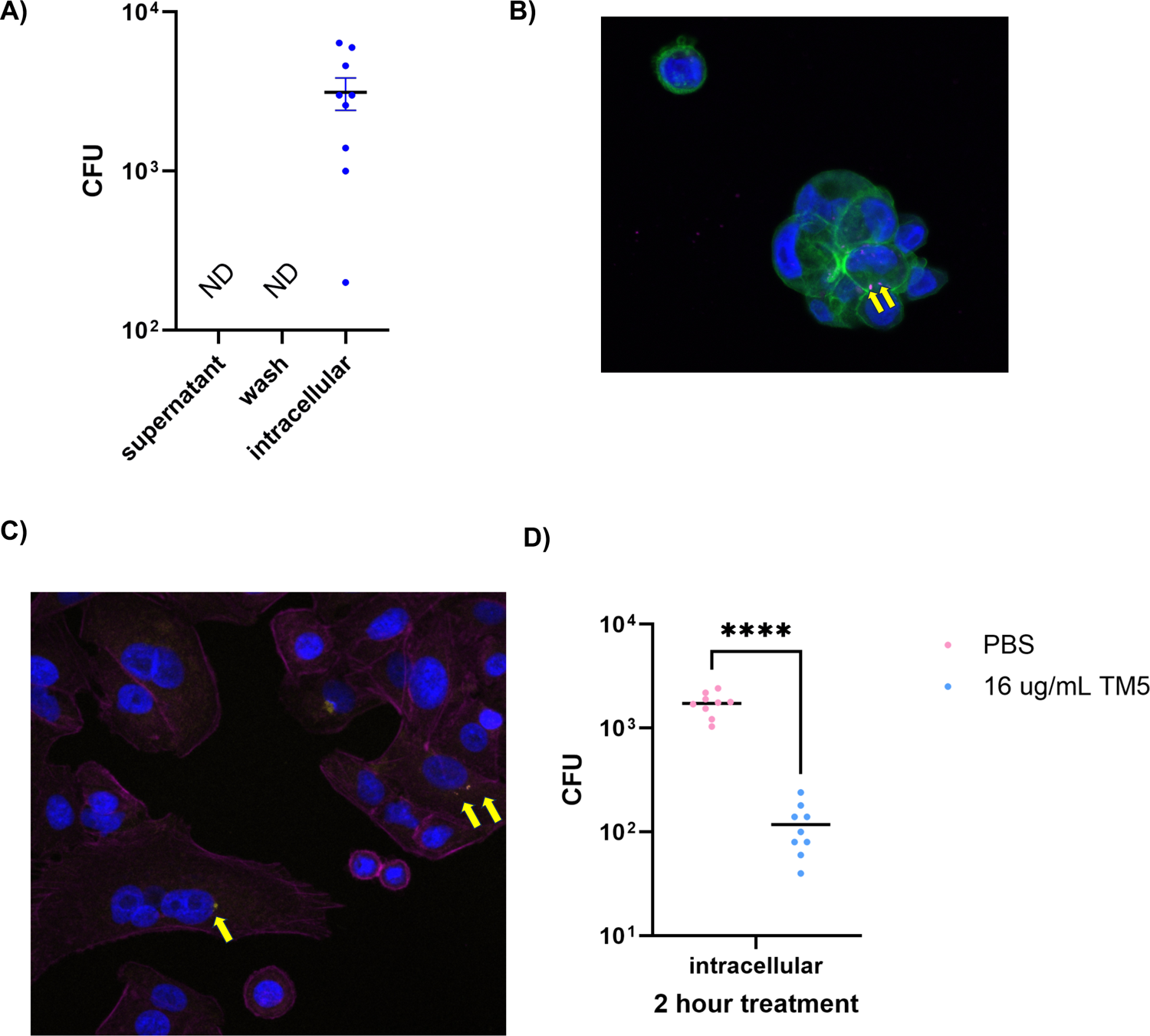
Establishment and treatment of intracellular PAO1 infection in Calu-3 lung epithelial cells a) A gentamicin protection assay was used to confirm intracellular PAO1 in Calu-3 cell, demonstrating that no bacterial CFU could be detected in the extracellular environment or following a wash step to remove adherent bacteria. In contrast, bacteria could consistently be detected in cell lysates. ND= not detected. b) Intracellular PAO1 within Calu-3 cells as visualized by confocal microscopy. Yellow arrows indicate individual bacteria stained with BacLight Deep Red. (Blue = DAPI, Green = Phalloidin) c) Intracellular PAO1 within A549 cells as visualized by confocal microscopy. Yellow arrows indicate individual bacteria stained with BacLight Deep Red. (Blue = DAPI, Green = Phalloidin) d) TM5 can treat intracellular PAO1 infections of Calu-3 cells at the nontoxic concentration of 16 μg/mL. A significant decrease (p<0.0001) in intracellular CFUs was detected following 2 hours of peptoid treatment as measured by Welch’s t-test.

### Treatment with Peptoid TM5 effectively kills intracellular *P. aeruginosa*

Having confirmed that *P. aeruginosa* PAO1 is able to establish an intracellular population within Calu-3 cells, we next evaluated whether peptoid TM5 was effective against intracellular PAO1. A concentration of 16 μg/mL of TM5 was chosen for therapeutic application as we have previously established this would be well tolerated by mammalian cells ^41^. Calu-3 cells were infected with PA01 as before, and following antibiotic treatment to eliminate extracellular bacteria the host cells were washed thoroughly to remove all traces of antibiotic. Fresh serum-free tissue culture media containing 16 μg/mL TM5 was added to each well and incubated at 37°C for two hours. Intracellular CFUs were then enumerated to evaluate the effect of peptoid treatment. After 2 hours, treatment with peptoid TM5 was able to significantly decrease the bacterial load of *P. aeruginosa* compared to untreated cells (Figure 2d). Host cells were monitored throughout the experiment and visually inspected to confirm that the decreased populations of bacteria were not due to death or detachment of the infected host cells. This confirmed that TM5 is effective at treating intracellular populations of bacteria, decreasing the intracellular bacterial load 10-fold after only a short incubation. The experiment was repeated three times on different days with *n=* 3 biological replicates per experiment.

### Identification of synergistic combinations of peptoid therapies

We next sought to further characterize the therapeutic potential of peptoid TM5 by evaluating its capacity for synergy with conventional antibiotics for use in combination therapies. Checkerboard assays are the most commonly used method for evaluating the effect of a combination of two therapeutics compared to the effect of the drug alone ^42^. Combinations of drugs can have different physiological effects, which can be quantified by the fractional inhibitory concentration index (FICI) value^43^. A combination of two drugs is synergistic when the FICI value is <0.5, which means that the MICs of both compounds has been decreased by at least 4-fold. A FICI value of <0.5-1 shows that the drug combination has an additive effect, resulting in a slight decrease in MIC value. An indifferent effect is observed when the FICI value is between 1 – 4, indicating the combination has no effect on the MIC. Sometimes, drugs can have a negative effect on each other, which is classed as antagonism when the FICI value is >4.

The antibiotics carbenicillin (CAR), chloramphenicol (CHL), kanamycin (KAN) and tetracycline TET) were selected for synergy testing due to their high MICs against clinical strains of *P. aeruginosa,* indicating resistance to these antibiotics (Table 1). These antibiotics are currently clinically irrelevant due to ineffectiveness against *P. aeruginosa* at approved clinical dosages or severe side effects in lung tissues at the MIC concentration^44, 45^. In addition, ceftazidime (CAZ), ciprofloxacin (CIP) and tobramycin (TOB) were chosen for evaluation as representatives of the current clinical gold standards against *P. aeruginosa* ^46^. Initial experiments were performed on both the laboratory and clinical strains, and subsequent confirmations were performed on the clinically relevant strain (Table 1).

**Table 1.** Confirmed synergistic combinations of antibiotic and peptoid TM5.

| Antibiotic | MIC antibiotic alone [ug/mL] | Synergistic concentration |  | FICI |
| --- | --- | --- | --- | --- |
|  |  | Antibiotic [ug/mL] | TM5 [ug/mL] |  |
| PA14 |  |  |  |  |
| Carbenicillin | 64 | 16 | 2 | 0.5 |
| Chloramphenicol | 128 | 2 | 4 | 0.5 |
| Tetracycline | 32 | 2 | 4 | 0.56 |
| PAZK2870 |  |  |  |  |
| Carbenicillin | 64 | 16 | 2 | 0.5 |
| Chloramphenicol | 128 | 64 | 0.5 | 0.31 |
|  |  | 32 | 2 | 0.5 |
|  |  | 16 | 2 | 0.38 |
|  |  | 1 | 4 | 0.5 |
| Tetracycline | 32 | 1 | 4 | 0.53 |
|  |  | 4 | 2 | 0.38 |

We observed potential synergy in our checkerboard assays with three of the antibiotics tested. Synergy of the combination of TM5 and tetracycline was observed against the clinical PAZK2870 strain, not however against the laboratory PA14 strain. Here the combination was additive, albeit one could argue that with a FICI value of 0.563 it was borderline additive/synergistic (Figure 3a). Carbenicillin showed synergy against all strains tested using a combination of 16 μg/mL Carb + 2 μg/mL TM5 which resulted in a FICI value of 0.5 (Figure 3b). Finally, chloramphenicol and TM5 worked synergistically against both PA14 and PAZK2870, at concentrations 32 μg/mL CHL + 2 μg/mL TM5 and 16 μg/mL CHL and 2 μg/mL TM5, with the FICI values being 0.31 and 0.38 respectively (Figure 3c).

**Figure 3.**
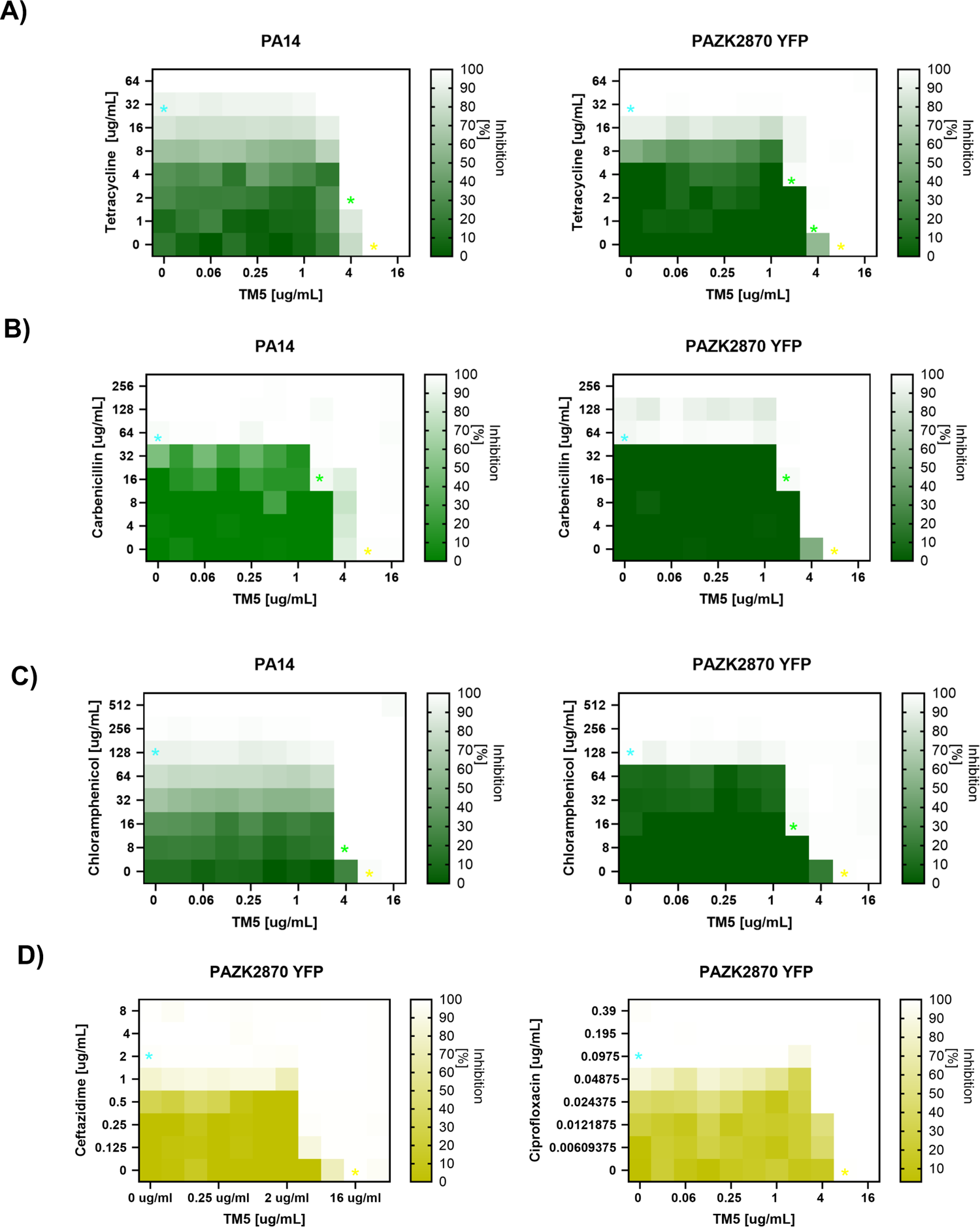
Synergistic combinations of TM5 with conventional antibiotics Checkerboard assays of TM5 in combination with a) tetracycline, b) carbenicillin and c) chloramphenicol indicated potential synergy with these antibiotics. 70 concentrations were examined per assay in single replicates– blue stars indicate MIC of antibiotic alone, yellow stars MIC of peptoid alone and a green star the synergistic combination. d) Checkerboard assays of TM5 and two antibiotics deemed additive (ceftazidime, ciprofloxacin)

The combination of kanamycin and TM5 resulted in a FICI value of 2.5 for PA14 and 1.5 for PAZK2870 and therefore, this combination was classed as indifferent. Similarly, tobramycin and TM5 was classified as indifferent with a FICI value of 2. Ciprofloxacin in combination with TM5 was exactly on the border of indifference and additive with a FICI value of 1. The combination of TM5 and carbenicillin was additive in both PAO1 and PAZK2870 with FICI values of 0.75. TM5 and ceftazidime had a FICI value of 0.625 and was therefore also additive (Figure 3d).

Checkerboard assays are designed to evaluate a range of combinations of different therapeutics, with only one replicate per combination. Therefore, we refined our analysis by confirming each synergistic combination using standard MIC assays performed in triplicate. Using this method, we confirmed that tetracycline (Figure 4a), carbenicillin (Figure 4b) and chloramphenicol (Figure 4c) all demonstrated synergy with TM5 and chose to move forward with a selected range of concentrations based on their FICI values. The concentrations of each antibiotic at which synergy was observed were below the MIC concentrations of antibiotic necessary to inhibit growth of *P. aeruginosa* PAO1 when used alone (Figure 4d), demonstrating rescue of the efficacy of these drugs when used in combination with peptoid TM5.

**Figure 4.**
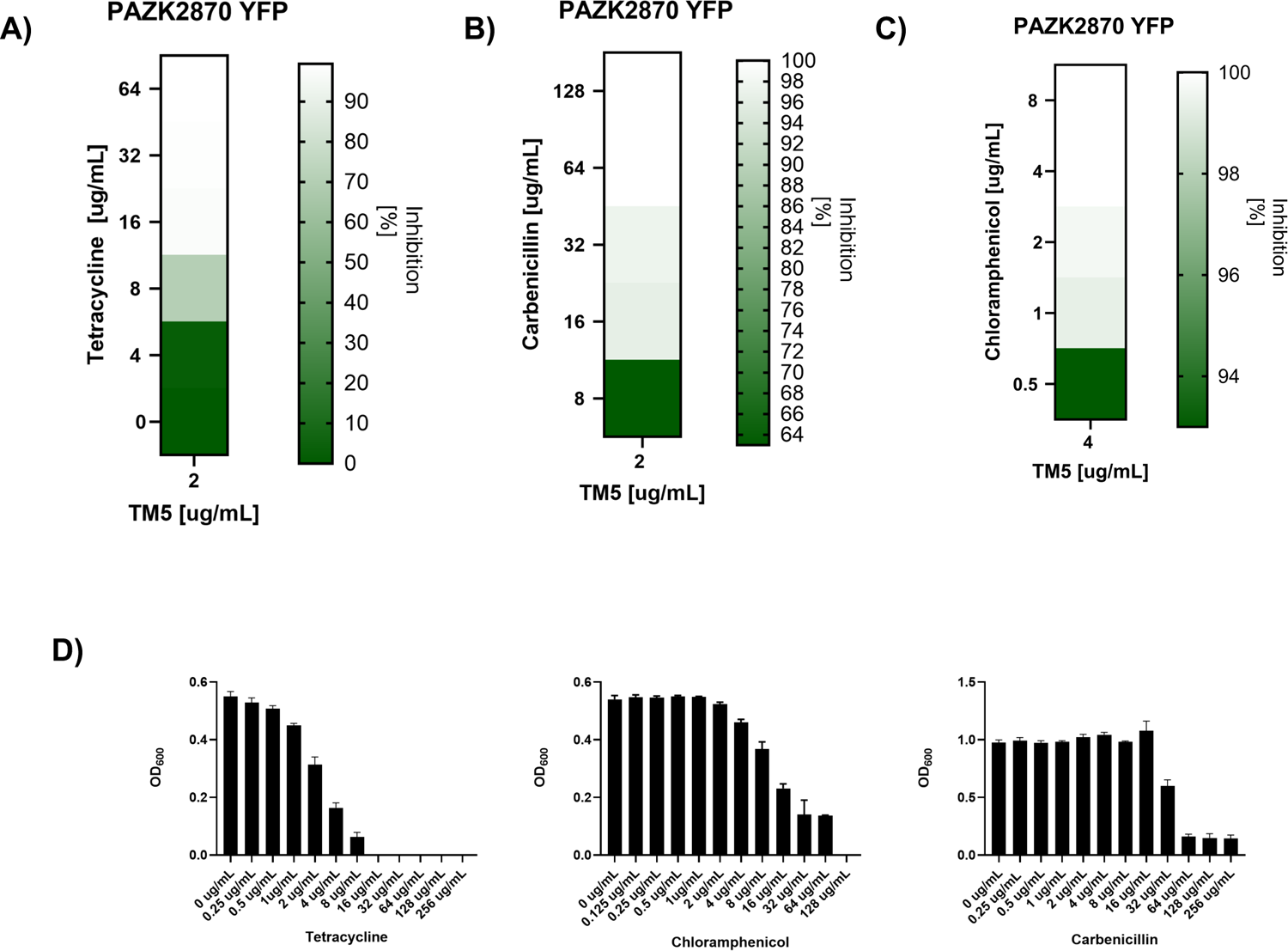
Confirmation of synergistic combinations of peptoid TM5 and antibiotics Each antibiotic with predicted synergy with TM5 was tested in triplicate using combined MIC assays to confirm the results of the checkerboard assays. a) Tetracycline is synergistic with 2 μg /mL TM5, b) carbenicillin is synergistic with 2 ug/mL TM5, and c) chloramphenicol is synergistic with a range of concentrations of TM5 up to 4 μg/mL. d) Individual MIC assays of the conventional antibiotics tetracycline, chloramphenicol and carbenicillin, showing the efficacy of each antibiotic alone. Graphs depict the mean of experiments performed in triplicate.

### Treatment of intracellular *P. aeruginosa* with synergistic combinations

We next evaluated the ability of selected synergistic combinations of peptoid and conventional antibiotic to eliminate intracellular *P. aeruginosa.* The combinations chosen for evaluation in Calu-3 cells were 16 μg/mL CAR + 2 μg/mL TM5, 16 μg/mL CHL +2 μg/mL TM5 and 4 μg/mL + 2 μg/mL TM5, based on their high synergy and low concentrations of each drug. Calu-3 cells were infected, washed, and treated for 2 hours with each synergistic combination as previously described, and bacterial CFUs were enumerated for each condition tested (Figure 5a,b). None of the combinations tested was as effective at eliminating intracellular populations of *P.* aeruginosa as TM5 alone at 16 μg/mL. However, we observed a significant decrease in intracellular CFU for the tested combinations of carbenicillin + TM5 and tetracycline + TM5, compared to no treatment. The combination of chloramphenicol and TM5 appears to be a borderline phenotype, showing a statistically significant in one independent experiment (Figure 5b), but only a non-significant downward trend in the other independent replicate (Figure 5a).

**Figure 5.**
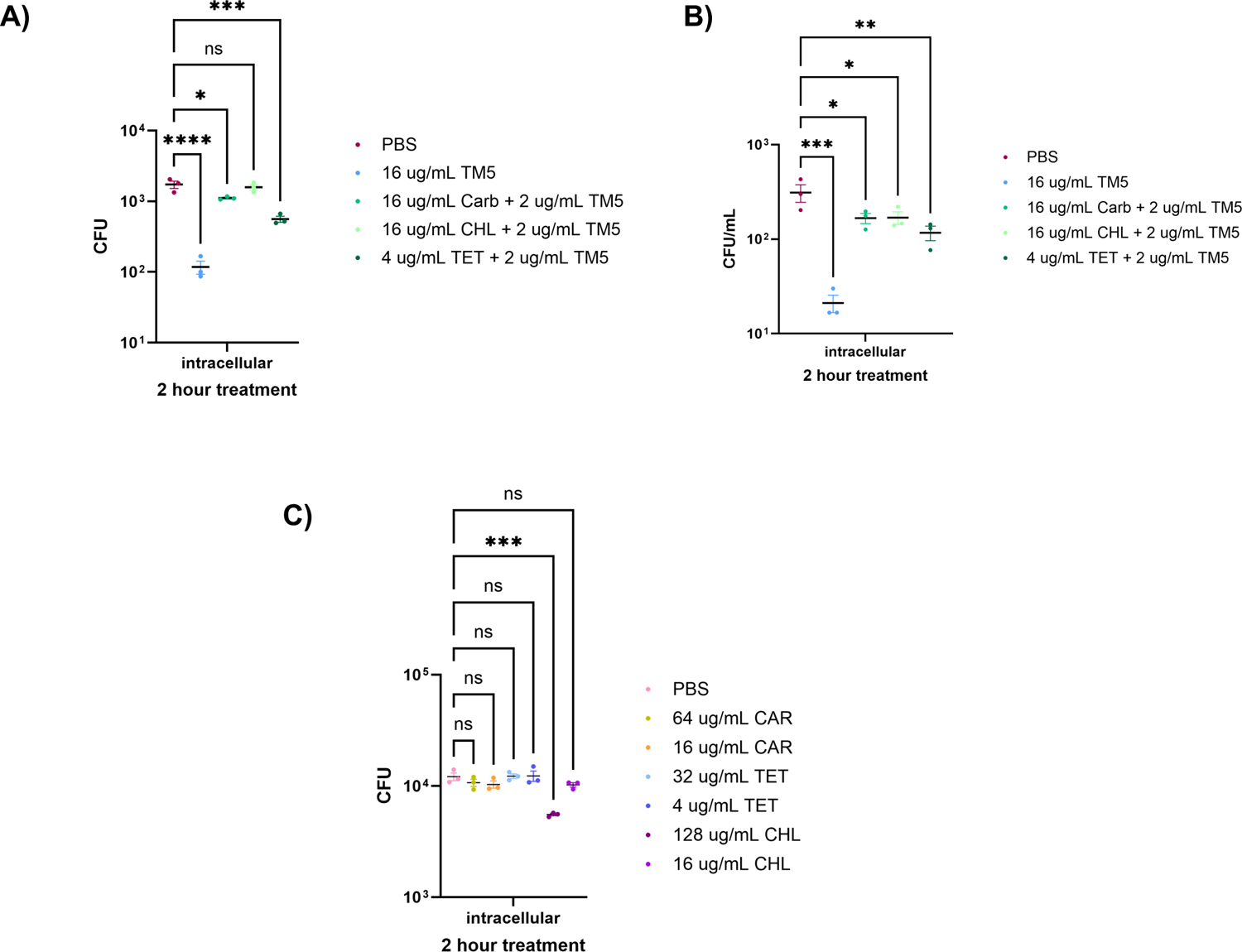
Synergistic antibiotic-peptoid combination treatment of intracellular *P. aeruginosa*. Calu-3 cells were infected with *P. aeruginosa* and treated with candidate synergistic concentrations of antibiotic and peptoid. There was no significant difference in intracellular CFU in cells treated with 16ug/mL CHL + 2ug/mL TM5 compared to untreated cells. For the other two combination treatments, 16 ug/mL CAR + 2 ug/mL TM5, and 4 ug/mL TET + 2 ug/mL TM5 a significant decrease in live intracellular bacteria was seen when compared to the infection not treated with antimicrobials. Two independent replicates are shown, each containing three biological replicates (a, b) Calu-3 cells treated with antibiotics individually at synergistic and MIC concentrations. None of the antibiotics were able to reduce intracellular CFUs at the concentrations used in synergistic therapies, while chloramphenicol was the only antibiotic that was able to decrease intracellular CFU at the much higher MIC concentration, potentially due to host cell toxicity at this concentration. Statistical testing was performed using ordinary one-way ANOVA tests (ns= not significant, * p≤0.05, **p≤0.01, *** p≤0.001, **** p≤0.0001)

To confirm that the ability of each combination therapy to eliminate intracellular bacteria was due to the inclusion of peptoid TM5, we also tested the efficacy of each antibiotic against intracellular bacteria individually. 1XMIC concentrations of each antibiotic were used to treat infected cells, including 64 μg/mL carbenicillin, 128 μg/mL chloramphenicol and 32 μg/mL tetracycline. Despite these concentrations being much higher than those used in the combination therapies, none of the conventional antibiotics were able to significantly reduce populations of intracellular bacteria alone, confirming that these antibiotics are unable to penetrate the host cell membrane at sufficient concentrations to eliminate bacteria within this niche (Figure 5c).

### Evaluating potential cytotoxic effects of synergistic peptoids treatment

A known problem with antimicrobial peptide-derived therapeutics is the potential for off-target cytotoxic effects on mammalian cells. Similarly, the conventional antibiotic chloramphenicol is notorious for its severe side effects at high concentrations. At the calculated *P. aeruginosa* MIC concentration of 128 μg/mL, this antibiotic would not be considered safe for use in human patients^45^. However, when used in combination with TM5, the concentration of CHL can be lowered to 16 μg/mL, well below the “safe” threshold of 25 μg/m^47^. However, it is important to evaluate the toxicity of each proposed combination to ensure the safety as well as the efficacy of treating an infection with both antimicrobials simultaneously.

To confirm that our combination therapies did not produce adverse effects, we assayed cell viability using two complementary colorimetric methods: (i) a resazurin assay which measures cell viability via the conversion of resazurin into the fluorescent dye resorufin and (ii) CellTiter 96® AQueous One Solution Cell Proliferation Assay, a variation of an MTT assay that measures cell metabolic activity through the reduction of a tetrazolium compound (MTS). Calu-3 cells were incubated with synergistic combinations and single concentrations of peptoid or antibiotic for 3 hours, and the percentage of viable cells was calculated following overnight incubation with resazurin or CellTiter solution. The viability of Calu-3 was found to remain unaffected by treatment with TM5, only demonstrating a slight decrease in viability at concentrations above 32μg/mL (Figure 6), well over the MIC. At the concentration used in our synergistic experiments (2 μg/mL), TM5 was extremely well tolerated by Calu-3 cells. Furthermore, no additional cytotoxicity was observed when the synergistic concentrations of CAR, CHL or TET were added to 2 μg/mL TM5, leading us to conclude that these combination therapies could be considered well tolerated and potentially candidates for future clinical therapies against chronic *P. aeruginosa* lung infections.

**Figure 6.**
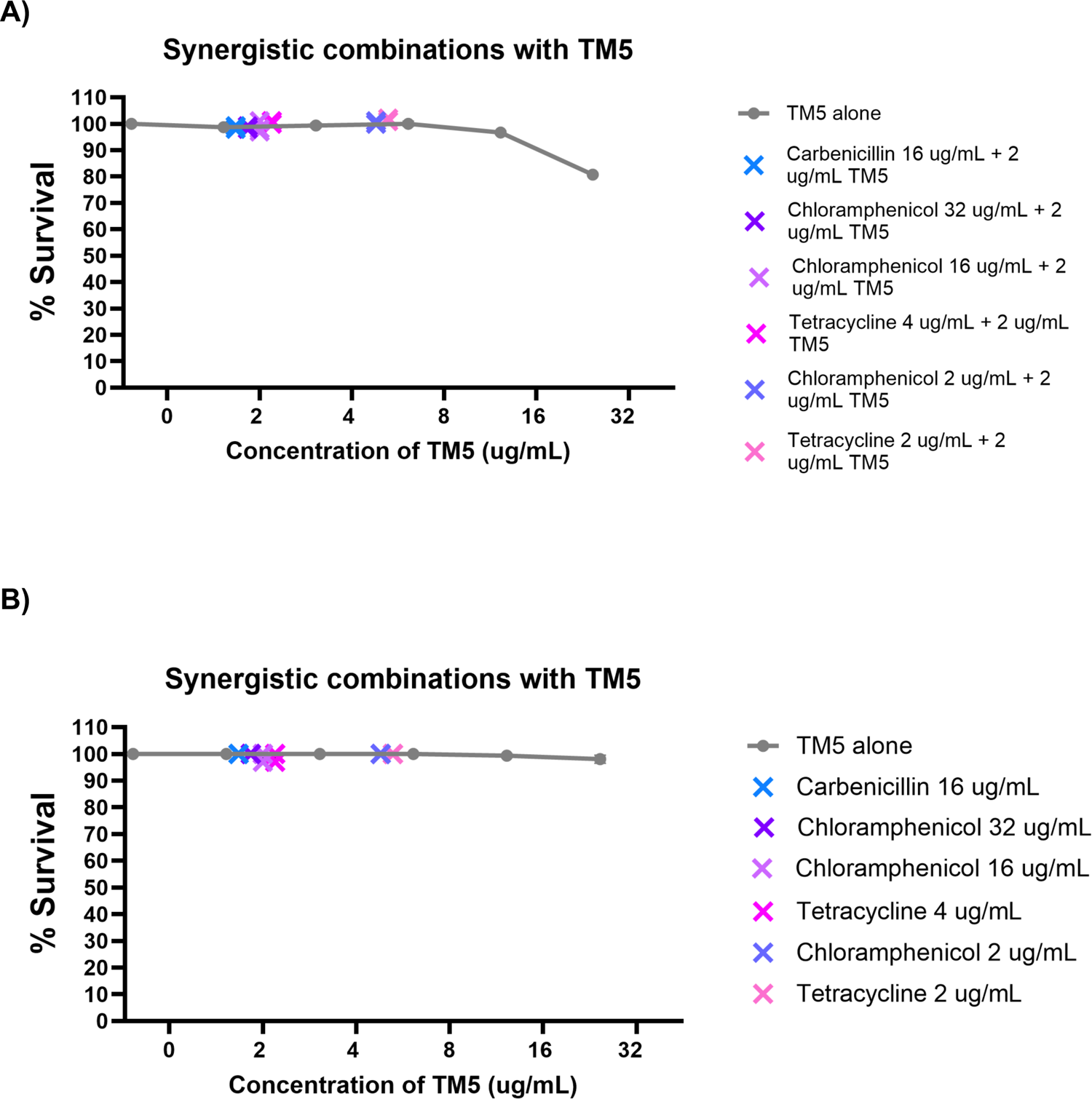
Cytotoxicity of TM5 and peptoid-antibiotic combinations in lung epithelial cells. Viability of mammalian cells following treatment with various antimicrobials was evaluated using resazurin (A) or MTS assay (B). TM5 treatment alone resulted in a slight decrease in survival of Calu-3 lung epithelial cells only at concentration of 32 ug/mL and above following 3 hour incubation with the peptoid. Various combinations of peptoid and antibiotic that demonstrated synergy (represented by Xs) were evaluated and found to result in no cytotoxicity to mammalian cells. Values shown represent means of experiments performed in triplicate.

## Discussion

Novel antimicrobials targeting gram-negative bacteria have become a rarity, while MDR cases continue to rise. Drug resistant infections have become a particular problem for highly vulnerable or immunocompromised patients, which leads to additional therapeutic challenges. Treatment strategies that can address chronic infections while avoiding further antimicrobial resistance development or severe adverse effects are therefore a research imperative. Here we suggest a novel antimicrobial peptoid as a treatment against bacterial populations associated with chronic infections, both as an individual therapy and in combination with conventional antibiotics. Such combination therapies represent a commonly used clinical strategy to attack infections from multiple angles, and have the potential to “rescue” drugs that have fallen out of clinical use due to antimicrobial resistance or toxicity.

We have previously shown that peptoid TM5 is a promising option to treat both respiratory tract and cutaneous infections with minimal cytotoxicity *in vivo*^30, 31^. In this study, we have further demonstrated the potential of TM5 to target intracellular bacteria, making this novel therapeutic a potential option for eliminating reservoirs of bacteria within host cells that are difficult to treat with conventional antibiotics. In addition, we have explored potential combination therapies utilizing this unique antimicrobial and evaluated the ability of these synergistic combinations to treat intracellular lung epithelial cell *P. aeruginosa* infections. While TM5 alone was most effective at eliminating intracellular bacteria, applying synergistic combinations may have additional therapeutic benefits.

Intracellular populations of *P. aeruginosa* have been proposed to represent an important bacterial reservoir in CF patients^48^. This intracellular niche could provide protection for the bacteria to evade the host immune system and could be an important factor in chronic infections. Persistent *P. aeruginosa* infections have been shown to persist after patients have been treated with both CFTR modulators and aggressive antibiotic regimes, and recurrent infections are a known clinical problem^49, 50^. A recent study demonstrated conclusive evidence of intracellular *P. aeruginosa* present in lung explant tissues from CF patients undergoing lung transplantation, providing strong evidence that intracellular reservoirs could be the source of chronic infections^14, 48^.

These intracellular niches likely contribute to the inability of current therapies to reliably eliminate chronic infections as many conventional antibiotics are not effective against intracellular bacteria^51^. Even antibiotics that are able to enter mammalian cells are generally present in sub-MIC concentrations intracellularly. This is problematic as exposure to subtherapeutic concentrations of antibiotics can drive bacterial expression of resistance and pro-biofilm genes^52^. In contrast, there is increasing evidence that peptide-based therapies are better able to overcome this barrier and target intracellular bacteria^53^. LL-37, the parental peptide of TM5, and other AMPs and their derivatives have been shown to be more effective than conventional antibiotics against intracellular *S. aureus*^54–56^. A synthetic AMP, OP145, has been shown to be effective against intracellular *Enterococcus faecalis* without harming host THP-1 macrophage cells, and the bovine-derived BSN-37 has been shown to be effective against intracellular *Salmonella*^57, 58^. Efficacy against intracellular *P. aeruginosa* has also been demonstrated for the synthetic peptides RR4 and RR4D in murine macrophages as well as the for the escualentin-1a-derived peptides Esc(1-21) and Esc(1-21)-1c in airway epithelial cells^59^. This growing realisation that peptide therapies have the potential to address intracellular reservoirs encouraged us to focus our studies on evaluating the potential of TM5 to eliminate intracellular populations of bacteria.

As *P. aeruginosa* has historically been studied as an extracellular pathogen and its intracellular lifecycle has not yet been fully elucidated, the development of a robust model of intracellular infection was an imperative first step for this study. To ascertain that we were indeed targeting intracellular bacteria and not adhered or planktonic populations, we adapted the traditional antibiotic protection assays commonly used to study intracellular pathogens to and thoroughly validated our model. Since *P. aeruginosa* demonstrates intrinsic resistance to many antibiotics, we evaluated different cell-impermeable antibiotics and tested a range of concentrations against our laboratory PAO1 strain. We determined that 1.5 hour treatment with 250 μg/mL of the antibiotic amikacin was sufficient to kill any bacteria adhered to the outer surface of the mammalian cells. Confocal microscopy was also used to further validate that we were in fact treating intracellular bacteria and provides further evidence of an intracellular stage of the *P. aeruginosa* life cycle.

With our newly established *in vitro* infection model, we were then able to demonstrate that TM5 can target and kill intracellular *P. aeruginosa*, decreasing the bacterial load 10-fold with even a short 2-hour treatment. However, our model does not current account for the thick mucous associated with CF lungs. The interference of mucous with efficacy will need to be tested in a more physiologically relevant model in future studies, such as air-liquid interface models, organoid models, or an *in vivo* mouse model of chronic *P. aeruginosa* infection that can mimic CF lung conditions^60, 61^.Intracellular accumulation already requires a higher concentration of antimicrobial than is necessary to treat planktonic bacteria, and this concentration would need to further increase to overcome the presence of thick CF mucous.

The exact mechanism by which TM5 is able to target intracellular bacteria is currently unclear, and future studies will be needed to determine whether or not TM5 is permeating mammalian membranes and directly killing *P. aeruginosa*. However, based on what is known about the parental AMP LL-37 and the lack of cytotoxic effects on mammalian cells we have observed, we can speculate that TM5 may have an immunostimulatory effect, initiating intracellular clearance indirectly by increasing bacterial killing by the host cells themselves rather than breaching the host membrane. This is of particular interest because combination therapies that combine immunostimulation and direct bacterial killing are a common practice in clinical settings, especially in patients with CF. These patients are rarely only dealing with one pathogen and have often been on long-term antibiotic treatment regimes or have developed chronic MDR infections. In standard CF treatment regimens, an antibiotic is often given in combination with an immunostimulatory antimicrobial, boosting the patient’s immune system to clear the infection and directly killing the bacteria^62^.

Tobramycin, an aminoglycoside, is often one of the first-line inhaled antibiotics administered in *P. aeruginosa* infections, and is commonly given in combination with the beta-lactam ceftazidime^63^. However, there are limitations to these first line treatment options. Aminoglycosides cannot penetrate mammalian cells and beta-lactams are often administered intravenously, which is associated with a higher risk for adverse effects including gastrointestinal issues, neurotoxicity and impaired renal function^23, 64^. Moreover, antimicrobial resistance has already been observed for all known treatment options, and the resistance rates against both aminoglycoside and beta-lactam antibiotics are only expected to rise.

Peptoid-based combination therapies have the potential to overcome many of the clinical challenges posed by current gold standard treatments including lack of cell permeability, adverse side effects, and antibiotic resistance. To the best of our knowledge, this work represents the first report of combination treatments of intracellular *P. aeruginosa,* as well as the first description of peptide synergy rescuing the efficacy of antibiotics which have been rendered obsolete through growing antimicrobial resistance. To identify these synergistic combinations, we employed checkerboard assays which allow for the quick and straightforward screening of multiple combinations of two antimicrobials at the same time^14^. FIC indexes are generally accepted as the best measure to provide a quantitative indication of interactions. However, the interpretation of these values is also often subject of debate and can vary between studies^65^. Checkerboard assays also do not provide any information on whether or not a combination is bactericidal as they only evaluate bacterial growth inhibition rather than viability.

We chose to focus on antibiotics that are no longer used to treat *P. aeruginosa* infections (carbenicillin), or to which *P. aeruginosa* is intrinsically resistant (chloramphenicol and tetracycline). CLSI (Clinical and Laboratory Standards Institute) are no longer maintained for carbenicillin, as most *P. aeruginosa* isolates are resistant to it^66^. We were able to confirm that both clinical and laboratory isolates of *P. aeruginosa* have MIC values for carbenicillin of at least 64 μg/mL, well over the resistance threshold (Table 1). However, combination therapy using our peptoid facilitated a reduction of the concentration of carbenicillin needed to inhibit bacterial growth down to 16 μg/mL, reopening the door for this once-useful antibiotic to return to clinical relevancy. When using chloramphenicol in combination with our peptoid the MIC could be lowered to 16μg/mL chloramphenicol which is below the toxic range of >25 μg/mL^45^. Interestingly, we did not see synergy of TM5 with aminoglycosides or current frontline antibiotics. This is most likely due to the fact that our strains do not show acquired resistance to these antibiotics and thus they are already highly effective at low concentrations. It would be of interest in future studies to perform checkerboard assays on strains that have naturally acquired resistance to these categories of drugs, such as tobramycin or ciprofloxacin. This would allow us to assess whether peptoid TM5 might be useful in rescuing the use of gold standard antibiotics against highly resistant strains.

Although, our combinations were less successful at eliminating intracellular bacteria than peptoid TM5 alone, combination therapies still hold potential benefits for side effect reduction as they allow the use of much lower concentrations of both drugs. In these studies, the effective concentration of TM5 is reduced from the MIC of 8 μg/mL down to 1-2 μg/mL, which greatly increases the therapeutic window of this new drug. However, it is also important to consider any potential impact on resistance emergence in combination therapies, as some previous studies have raised concerns that treatment with multiple drugs could potentially select for new resistance emergence, while others have provided evidence that combination therapy can delay the evolution of multi-drug resistance^67–69^. As such, it is important to evaluate every synergistic combination carefully and consider factors such as mechanism of action and any potential for broad-spectrum collateral sensitivity^70, 71^. As the proposed mechanism of TM5 is the destabilisation of the bacterial membrane, something that cannot be overcome through mutation of a single gene, the evolution of resistance to TM5 is less likely. We have confirmed this in laboratory evolution experiments, and believe this makes TM5 a particularly good candidate for combination therapy^31, 35^. Moreover, synergistic combination therapies are overwhelmingly more likely to result in full elimination of an infection and more positive clinical outcomes. Combination therapies consistently outperforming single drug therapies, suggesting that the benefits of combinatorial treatment are likely to outweigh any potential risks^72, 73^.

In conclusion, this work represents an exciting step forward towards the development of novel therapeutic options against the chronic *P. aeruginosa* infections often associated with cystic fibrosis. Peptoid TM5 appears to have the ability to target intracellular populations of bacteria, and future studies will focus on elucidating the mechanism through which this peptoid is able to accomplish this, whether via direct penetration of mammalian cells or immunostimulation to enhance the ability of host cells to clear *P. aeruginosa* infections. Moreover, we have shown that TM5 shows great promise as a combination therapy due to the synergy observed with conventional antibiotics that have lost the potential for clinical use individually. Based on our previous studies evaluating the safety of TM5 in an *in vivo* model, we believe that the extremely low concentrations of each drug able to successfully clear intracellular *P. aeruginosa* in combination therapies will be well-tolerated clinically. Future studies will focus on evaluating these novel therapies *in vivo*, and determining if these combinations are similarly effective against other bacterial pathogens that exhibit intracellular bacterial reservoirs^31^.

## Methods

### Peptoid Design and Synthesis

Peptoid synthesis was performed at the Molecular Foundry in the Lawrence Berkeley National Laboratory, Berkeley, CA using a Symphony X (Gyros Protein Technologies) as has been previously described^31, 35^. In brief, a Rink amide MBHA resin (EMD Biosciences) was used to prepare the peptoids, followed by rifluoroacetic acid (TFA):triisopropylsilane:water (95:2.5:2.5 volume ratio) treatment for 10 minutes to remove synthesized peptoids from the MBHA resin. Purification was then performed using a A C18 column in a reversed-phase high performance liquid chromatography (HPLC) system (Waters Corporation), employing a linear acetonitrile and water gradient with a compound purity greater than 95%^74^. Successful peptoids synthesis was confirmed via electrospray ionization mass spectrometry.

### Bacterial Strains and Culture Conditions

Laboratory *P. aeruginosa* strains PAO1 and PA14 were used as reference strains and *P. aeruginosa* strains PAZK2019, PAZK69, PAZK3095, and PAZK2870 from the Kolter collection (Oxford) were used as clinically relevant strains for peptoid efficacy testing. Cation-adjusted Mueller-Hinton (MH) agar and broth (BD Biosciences) was used for all peptoid experiments, and Luria-Bertani (LB) broth and agar (Fisher) were used for overnight cultures and CFU enumeration. All bacterial growth was performed at 37 °C with, with liquid cultures inoculated from individual colonies and incubated at 37 °C with shaking at 250 rpm in a Solaris 4000I Incubated Shaker.

### Minimum Inhibitory Concentration (MIC) and Minimum Bactericidal Concentration (MBC) Assays

A standardised procedure was used to determine the MICs of chosen the chosen antibiotics: carbenicillin (CAR), ceftazidime (CAZ), chloramphenicol (CHL), ciprofloxacin (CIP), kanamycin (KAN), tetracycline (TET), tobramycin (TOB), and peptoid TM5. Briefly, 2-fold dilutions of each antibiotic, peptoids, or combination therapy were prepared in MH broth and introduced into polypropylene 96-well plates (Greiner) win triplicate. An overnight culture of each *P. aeruginosa* strain was then diluted 1:100 and regrown to log phase (OD_600_ between 0.4 – 0.9) shaking at 37 °C to ensure that bacteria were in an exponential growth phase. These log phase cultures were diluted to OD_600_ 0.001 and added individually to each well of the plate. Plates were then incubated overnight at 37 °C. For each plate, bacterial cultures without antimicrobial were prepared as a positive control, and MH broth without antibiotic served as a negative control. The following day, each well was gently resuspended and 100uL of each well was transferred to a polystyrene plate. OD_600_ readings were then taking on a FluoSTAR Omega plate reader. MIC was defined by a 90% decrease in OD_600_ compared to untreated bacteria. MBC assays were performed using the same protocol, with the exception that following the overnight incubation an aliquot of each sample was taken and 10-fold dilutions were prepared in a new 96-well plate. Each sample was then spotted onto MH agar plates and incubated overnight at 37°C and bacterial colony forming units (CFUs) were enumerated the next day. All experiments were performed in triplicate with a minimum of two independent replicates.

### Checkerboard assays

Checkboard assays were setup in 96-23ll polypropylene plates (Greiner) using various antibiotics (drug A) and peptoids TM5 (drug B). A stock solution of drug A at a concentration 4-fold higher than the highest concentration to be tested was prepared in MH broth. A separate stock solution of drug A was prepared at a concentration 8-fold higher than the highest concentration to be tested, which was chosen based on the MICs of each antimicrobial. A 2-fold dilution of drug A was then introduced into each well from row A downwards. Wells 1A to 10A contained the 4x solution and well 12A contained the 8x solution. A stock solution of drug B was obtained by preparing a solution 4-fold of the highest concentration to be tested. This was then dispersed into wells 11A – 11G and a 2-fold dilution was performed to column 2. The plate was incubated overnight at 37 °C. On the next day, the wells were resuspended and 100uL was transferred to a polystyrene plate in order to read the OD600 via a plate reader. The MIC of each drug individually was established - for drug a 1A – 1G, for drug B 11H-2H and used to evaluate the FICI values of the combinations. The following calculation was used to determine the FIC value for each peptoid-antibiotic combination:

The combination was then classed as synergistic (FICI <0.5), additive (FICI value of <0.5-1), indifferent (FICI = 1-4) or antagonistic (FICI>4) according to the FIC value.

### Calu-3 and A549 infection with *P. aeruginosa* strains

Calu-3 cells were growth in Minimal Essential Medium Eagle (Gibco) supplemented with 2 mM L-Glutamine (Gibco), 10 mM HEPES (Gibco) and 10% FBS (Fisher).

A549 cells were grown in Dulbecco’s Modified Eagle’s Medium (DMEM) supplemented with 10% FBS. Cells were seeded the day before infection at 2 × 10^5^ cells per well in a 24-well tissue culture plate (Corning). The following day, the cells were infected with bacterial cultures grown to log phase and diluted to MOI 1 (2 × 10^5^ CFU/well) in a 1mL volume of MEM. The cells were incubated with bacteria for 1 hour at 37°C and then washed with PBS. Extracellular bacteria were killed with fresh media containing 250μg/mL amikacin (Sigma), incubated for 1.5 hours. The cells were then again washed with PBS. For treatment conditions, 16μg/mL TM5 or synergistic peptoid-antibiotic concentrations were applied for 2 hours. Calu-3 cells were then lysed with 0.1% Triton-X, and 10-fold dilutions of each sampled were spot plated on LB or MH agar in triplicate to enumerate intracellular CFUs.

### Cytotoxicity assays

Cytotoxicity assays were performed using CellTiter 96® AQueous One Solution Cell Proliferation Assay kit (Promega) or resazurin assay as has been previously described^75^. Calu-3 cells were seeded into 96-well plates at 1×10^4^ cells/well. The following day, antibiotic, peptoid or antibiotic-peptoid combinations were diluted in cell culture media without FBS at desired concentrations and incubated at 37°C for 3 hours. 20uL of the CellTiter 96® AQueous One Solution Reagent was then added to each well and incubated for another 4 hours, and the plate was then read at absorbance 490nm.

### Immunofluorescence Staining and Microscopy

Calu-3 or A549 cells were seeded onto glass cover slips in 24-well plates at 2 × 10^5^ cells per well in and infected as described above with *P. aeruginosa* PAO1 that had been stained with BacLight Red (Invitrogen) following the manufacturer’s protocol and the mammalian cells were infected with labelled bacteria for 1 hour with a MOI of 10. Antibiotic protection assays was performed as before to eradicate all extracellular bacteria. After an additional 1.5 hours, the media was removed, each well was washed with PBS, and the coverslips were then fixed with 4% paraformaldehyde (PFA) for 20 minutes. Following fixation, the PFA removed and the cells were washed with PBS, permeabilised in 0.1% Triton-X for 5 minutes at room temperature and blocked with 1% BSA in PBS for a minimum of one hours. The coverslips were then stained with Phalloidin and DAPI (Invitrogen) for one hour, washed 3X with PBS, and .mounted onto glass slides using Vectashield (VectorLabs) mounting media. Imaging was performed on a Zeiss LSM 880 (Airyscan) at the University of Edinburgh Light Microscopy Core.

## Acknowledgements

AEB thanks the NIH for funding this work with a Director’s Pioneer Award, grant # 1DP1 OD029517. AEB also acknowledges funding from SENS Research Foundation (now Lifespan Research Institute), Stanford University’s Discovery Innovation Fund, the Cisco University Research Program Fund, the Silicon Valley Community Foundation, and Dr. James J. Truchard and the Truchard Foundation. We gratefully acknowledge Dr. Michael Connolly and Dr. Behzad Rad at the Molecular Foundry for assistance with peptoid synthesis and sample preparation equipment. Work at the Molecular Foundry was supported by the Office of Science, Office of Basic Energy Sciences, of the U.S. Department of Energy under Contract No. DE-AC02-05CH11231. MM and MB thank the UKRI Medical Research Council and the University of Edinburgh for funding to support this work through the Precision Medicine Doctoral Training Programme. We acknowledge Dr. Joanne Thompson, Dr. Simon Martin, Dr. Jason Mooney and Dr. Thamarai Schneiders for their discussions and feedback on the project.

